# Sex-dependent development of *Kras*-induced anal squamous cell carcinoma in mice

**DOI:** 10.1101/2021.08.04.455138

**Authors:** Morgan T. Walcheck, Kristina A. Matkowskyj, Anne Turco, Simon Blaine-Sauer, Manabu Nukaya, Jessica Noel, Oline K. Ronnekleiv, Sean M. Ronnekleiv-Kelly

## Abstract

Anal squamous cell carcinoma (SCC) will be diagnosed in an estimated 9,080 adults in the United States this year, and rates have been rising over the last several decades. Most people that develop anal SCC have associated human papillomavirus (HPV) infection (∼85-95%), with approximately 5-15% of anal SCC cases occurring in HPV-negative patients from unknown etiology. This study identified and characterized a *Kras*-driven, female sex hormone-dependent development of anal squamous cell carcinoma (SCC) in the *LSL-Kras*^*G12D*^ ; *Pdx1-Cre (KC)* mouse model that is not dependent on papillomavirus infection. One hundred percent of female KC mice develop anal SCC, while no male KC mice develop tumors. Both male and female KC anal tissue express Pdx1 and Cre-recombinase mRNA, and the activated mutant *Kras*^*G12D*^ gene. Although the driver gene mutation *Kras*^*G12D*^ is present in anus of both sexes, only female KC mice develop *Kras*-mutant induced anal SCC. To understand the sex-dependent differences, KC male mice were castrated and KC female mice were ovariectomized. Castrated KC males displayed an unchanged phenotype with no anal tumor formation. In contrast, ovariectomized KC females demonstrated a marked reduction in anal SCC development, with only 15% developing anal SCC. Finally, exogenous administration of estrogen rescued the tumor development in ovariectomized KC female mice and induced tumor development in castrated KC males. These results confirm that the anal SCC is estrogen mediated. The delineation of the role of female sex hormones in mediating mutant *Kras* to drive anal SCC pathogenesis highlights a subtype of anal SCC that is independent of papillomavirus infection. These findings may have clinical applicability for the papillomavirus-negative subset of anal SCC patients that typically respond poorly to standard of care chemoradiation.

## Introduction

In 2021, an estimated 9,080 adults will be diagnosed with anal squamous cell carcinoma (SCC) in the United States, and anal SCC has been increasing in incidence over the last several decades.[1,2] Anal SCC is typically associated with human papillomavirus (HPV) infection (∼85-95%),[2,3] yet approximately 5-15% of anal SCC cases occur in HPV-negative patients with unknown etiology.[4,5] Unfortunately, patients with HPV-negative anal SCC are significantly less responsive to standard of care chemoradiation[5], and have a worse prognosis than HPV-positive anal SCC.[6] This study presents a novel etiology for HPV-negative anal SCC development driven by mutant *Kras*.

In human anal cancer, mutant *Kras* is identified in 10% of HPV-negative anal SCC samples.[7] Despite this association, to our knowledge, the present study is the first to identify this correlation in a pre-clinical model. The mutant *Kras*-driven development of anal SCC was detected in a genetically engineered mouse model (GEMM) traditionally used in the investigation of pancreatic ductal adenocarcinoma (PDAC). This mouse harbors a *Kras*-mutation (*Kras* ^*G12D*^) in cells expressing Cre-recombinase from pancreatic and duodenal homeobox 1 (*Pdx1*) promoter (KC mice: *Lox-stop-lox Kras*^*G12D/+*^ ; *Pdx1-Cre*)[8]. In this study, we found that Pdx1 expression and consequent Cre-recombinase expression in the anal epithelium caused activation of the *Kras*^*G12D*^ gene in the anal epithelium and tumor development. Further, we observed that only female mice developed anal SCC suggesting a sex-hormone dependent interaction with *Kras* ^*G12D*^ that triggers tumor formation.

Therefore, we sought to understand the sex-dependent development of anal SCC in KC mice. Activated *Kras*^*G12D*^ was present in the anal tissue of both sexes of KC mice, suggesting both sexes have the potential to develop *Kras*-mutant anal SCC. To ascertain why only female KC mice develop tumors, we ovariectomized females and castrated males to eliminate endogenous sex hormones production in the mice and found ovariectomized females displayed significantly reduced anal tumor development, signifying female sex hormone dependence. In turn, ovariectomized and castrated KC mice dosed with estrogen resulted in tumor development in both KC female and KC male mice, respectively, indicating the anal tumor development is estrogen mediated. This novel phenotype shows a female sex hormone dependent pathogenesis of *Kras*-mutant anal SCC that is independent of HPV infection. Given that 2-5% of anal SCC overall and 10% of HPV negative anal SCC harbor *Kras*-mutations [7], these findings may have therapeutic implications for this subset of patients. Lastly, the sex-based difference highlights the importance of characterizing both sexes in pre-clinical studies.

## Methods

### Animals

All animal studies were conducted according to an approved protocol (M005959) by the University of Wisconsin School of Medicine and Public Health (UW SMPH) Institutional Animal Care and Use Committee (IACUC). Mice were housed in an Assessment and Accreditation of Laboratory Animal Care (AALAC) accredited selective pathogen-free facility (UW Medical Sciences Center) on corncob bedding with chow diet (Mouse diet 9F 5020; PMI Nutrition International), and water *ad libitum*. The *Lox-Stop-Lox (LSL) Kras*^*G12D*^ (B6.129S4-Kras ^tm4Tyj^/J #008179), *Pdx1-Cre* (B6.FVB-Tg(*Pdx1-cre*)^6Tuv^/J) and Ai14 (B6.Cg-Gt(ROSA)26Sor^tm14(CAG-tdTomato)Hze^/J #007914) mice were purchased from the Jackson Laboratory (Bar Harbor, ME) and housed under identical conditions. All mice listed are congenic on a C57BL/6J background (backcrossing > 15 generations). The *LSLKras*^*G12D*^ and *Pdx1-Cre* mice were bred to develop *LSL*-*Kras*^*G12D*^ ; *Pdx1-Cre* (KC) mice. The Ai14, *Pdx1-Cre* and *LSLKras*^*G12D*^ were bred to develop *Rosa26*^*LSL-tdTomato*^ ; *LSLKras*^*G12D*^ ; *Pdx1-Cre* (AiKC) mice. Genotyping was performed according to Jackson Laboratory’s protocols (Cre: Protocol #21298, Kras: Protocol #29388 and Ai14: Protocol #29436). Original observations were performed in 16 male KC and 14 female KC mice and 83 control genotypes. Based on stark sex-dependence, we calculated that 10 KC male, 10 KC female, and 12 control mice were needed for castration / ovariectomy studies to detect a 50% change in tumor formation by Fisher Exact test and alpha of 0.05. We then calculated that 12 KC female mice would be sufficient for the E2 dosing studies (6 E2 dosed and 6 sham controls) as well as 12 KC male mice (6 E2 dosed and 6 sham controls). Finally, we used 6 AiKC mice to visually confirm the location of Pdx1-Cre (projected location of mutant Kras expression) in the anal tissue. The health and well-being of the mice were monitored closely by research and veterinary staff. Mice that showed signs of distress such as disheveled coat, hunched posture, rapid weight loss, lack of feeding or lack of defecation were immediately euthanized. During the experiment process, one castrated KC male mouse was euthanized due to decline in health and one castrated KC male mouse, one E2 dosed ovariectomized KC female, one sham dosed castrated KC male and two E2 dosed castrated KC male mice were found deceased of uncertain circumstances before the study end point. These mice were not included in the results. Mice were euthanized through CO^2^ asphyxiation.

### Genotyping for the activation of *Kras*^*G12D*^ mutation construct

Activated *Kras*^*G12D*^ refers to the successful Cre-mediated excision of the *Lox-Stop* sequence, allowing for transcription of the mutant *Kras* allele. To determine the tissue specific activation of the *Kras*^*G12D*^ mutation, we followed the standard method first published by Hingorani [8,9] and further utilized by other groups working with this *Lox–Stop–Lox* conditional *Kras* mouse strain[10–12]. Genomic DNA was isolated from tail, pancreas, anus and anal tumor from KC mice. The DNA was then amplified using polymerase chain reaction (PCR) with the following probes: 5’-GGGTAGGTGTTGGGATAGCTG-3’ (OL8403) and 5’-CCGAATTCAGTGACTACAGATGTACAGAG-3’ (OL8404) with conditions previously published[11]. These primers amplified a 325 bp band corresponding to the activated *Kras*^*G12D*^ mutant allele and a 285 bp band corresponding to the WT allele.

### Tumor studies

The study endpoint (age 9 months) was selected based on existing data evaluating and reporting on male KC mice at this age[8]. At 9 months, mice were euthanized and underwent cervical dislocation followed by midline laparotomy for solid organ assessment. The anus was also removed for pathologic analysis. A board-certified surgical pathologist with subspecialty training in gastrointestinal pathology (KAM) who was blinded to the mouse genotype and sex evaluated the histologic sections.

### Histology

KC mouse tissues (anus and pancreas) were fixed in 10% buffered formalin for 48 hours. Serial 4 μm sections from paraffin-embedded tissues were mounted on charged slides. Hematoxylin and eosin (H&E) was performed by the Experimental Animal Pathology Lab (EAPL) at the University of Wisconsin-Madison. The histology was evaluated by a certified pathologist (KAM).

### DNA recovery from H&E stained formalin fixed paraffin embedded (FFPE) samples

The anal tissue was scraped from H&E stained slides using a sterile blade.[13] The deparaffinization and genome DNA extraction from H&E stained anal tissues was performed according to manufacturer’s instructions using ReliaPrep FFPE gDNA MiniPrep System (Promega, Madison, WI).

### Tdtomato immunohistochemistry (IHC)

IHC staining for red fluorescence protein (tdTomato) was performed by the Experimental Animal Pathology Lab (EAPL) at the University of Wisconsin-Madison. For IHC staining, sections were deparaffinized in xylenes and hydrated through graded alcohols to distilled water. Antigens were retrieved using citrate buffer pH 6.0 (10 mM Citric Acid, 0.05% tween 20). Endogenous peroxidase was blocked with 0.3% H_2_O_2_ in PBS for 10 minutes at room temperature and blocking of non-specific binding was performed using 10% goat serum. Sections were incubated with rabbit anti-RFP antibody (600-401-379, Rockland Inc, Pottstown, PA) (1:1600) overnight at 4ºC. After rinsing, sections were incubated with ImmPRESS goat anti-rabbit IgG (MP-7451, Vector Laboratories, Burlingame, CA) for 30 minutes at room temperature. Detection was performed using DAB substrate kit (8059S, Cell Signaling Technology, Danvers, MA). Samples were counterstained using Mayer’s hematoxylin (MHS32, Millipore-Sigma, St. Louis, MO) for one minute.

### RNAScope in situ hybridization

MmuPV1 detection was performed using the RNAscope 2.5 HD Assay-Brown kit (Advanced Cell Diagnostics, Newark, CA; 322300) and probe to MmuPV1 E4 (473281) as previously described.[14] NSG mouse anal tissues that were infected with MmuPV1 or mock infected [14] were included as positive and negative controls, respectively.

### Estrogen receptor alpha immunofluorescence

Formalin-fixed (10 % formalin), paraffin-imbedded mouse tissue sections mounted on Superfrost Plus glass slides (Fisher Scientific, Pittsburgh, PA), were deparaffinized with Xylene (3 x 5 min), and rehydrated in descending concentrations of ethanol as follows: 2 x 10 min each in 100%, 95%, 70%, and 50% ethanol followed by two washes in deionized water for 5 min each and a final wash in phosphate buffer (PB; 0.1 M phosphate buffer, pH 7.4) solution for 10-15 min. Sections were pretreated with normal donkey serum solution (3% donkey serum, 0.3% Triton-X 100 in PB, pH 7.4) for 30 min at room temperature and then washed briefly in PB before being incubated for 48 hrs at 4° C with an estrogen receptor α (ERα) rabbit antibody (C1355) diluted 1:5000. Thereafter the sections were rinsed in PB and next incubated with biotinylated donkey anti-rabbit IgG (1:500) for 2 hours at room temperature, another wash for 30 min in PB and then reacted with streptavidin Alexa Fluor 594 (1:2500) for 3 hrs. Both primary and secondary antisera were diluted in Tris-(hydroxymethyl)aminomethane (0.5%; Sigma-Aldrich) in phosphate buffer containing 0.7% seaweed gelatin (Sigma-Aldrich), 0.5% Triton X-100 (Sigma-Aldrich), and 3% BSA (Sigma-Aldrich), adjusted to pH 7.6. Adjacent sections were treated equally, but without the ERα antibody for control purposes. After a final rinse overnight in PB, the sections were cover-slipped with gelvatol containing the anti-fading agent 1,4-diazabicyclo(2,2)octane (DABCO; Sigma-Aldrich; 50 mg/ml). Sections were screened and photographed using a Nikon E800 fluorescent microscope (Eclipse E800; Nikon Instruments, Melville, NY) equipped with a fiber illuminator (Intensilight C-HGFI; Nikon Instruments) and a high-definition digital microscope camera head (DS-Fi1; Nikon Instruments) interfaced with a PC-based camera controller (DS-U3; Nikon Instruments). It should be noted that the C1355 ERα antibody has been documented to be specific for ERα in rat and mouse tissues and does not recognize ERβ.[15]

### DNA recovery from FFPE tissues and MmuPV1 detection by PCR

DNA was isolated from two formalin fixed paraffin embedded slides per sample as previously described.[14] PCR was performed using primers specific to the MmuPV1 genome in the L1 region (F: 5’-GGAAGGAGAGAGCAAGTGTATG-3’, R: 5’-GGGTTTGGTGTGTTGGTTTG-3’) and analyzed via agarose gel.

### RNA isolation

Immediately following cervical dislocation and resection of the organs, specimens (pancreas and anus) allocated for RNA isolation were placed into RNAlater (ThermoFisher Scientific, Waltham, MA). The RNA isolation commenced immediately using Qiazol lysis and homogenization using a tissue homogenizer (Brinkmann Instruments, Model PT 10/35, 110 Volts, 6 Amps, 60 Hz). RNA was isolated using the Qiagen RNeasy Kit (Qiagen, Hilden, Germany). The extracted RNA was quantified using a spectrophotometer (ClarioStar Plate Reader, BMG LABTECH, Ortenberg Germany) and diluted to 50 ng/μL. Electrophoresis of the purified RNA was performed with the Agilent 2100 Bioanalyzer (Agilent, Santa Clara, CA), and each sample demonstrated an RNA Integrity Number (RIN) of 7.5 or higher, indicative of high-quality RNA.

### Quantitative reverse transcription PCR

The qPCR was done as previously described.[16] Briefly, 500 ng of RNA was reverse transcribed using the High Capacity cDNA Reverse Transcription Kit (Thermo Fisher, Waltham, Ma) per manufacturer protocol. The qPCR was performed on the Thermo Fisher QuantStudio 7 (Thermo Fisher, Waltham, Ma). All reactions were run in triplicate. Results were analyzed using the delta-delta CT method. Expression levels were calculated relative to the average of the C57BL6/J female mice (baseline) or the average of the KC females. The reference group was labeled on each graph. The following TaqMan® probes were used: *Cre* (Enterobacteria phage P1 cyclization recombinase, Mr00635245_cn), *Pdx1* (pancreatic and duodenal homeobox 1, Mm00565835_cn) and the house keeping gene *Hprt* (hypoxanthine guanine phosphoribosyl transferase, Mm03024075_m1) (Thermo Fisher, Waltham, Ma).

### Castration and ovariectomy

To evaluate the sex differences of anal SCC development in KC mice, male WT and male KC mice were castrated; meanwhile, female WT and female KC mice were ovariectomized at 6-7 weeks of age. Mice were anesthetized with isoflurane inhalation throughout the surgery. Slow-release buprenorphine was used as an analgesic for mice undergoing this surgery. The hair from the surgical area was removed with clippers, and the surgical area was sterilized with an iodine scrub. Under sterile conditions and using sterilized tools, the testis and ovaries were removed from male and female mice, respectively[17,18]. To remove the testis, gentle pressure was applied to the abdomen to push the testis into the scrotal sac [17]. A short 10mm midline incision was made through the skin in the middle of the abdomen [17]. The testis were located, gently pulled out through that incision along with the epididymal fat pad and carefully removed via cauterization [17]. To remove the ovaries, two incisions were made: short dorsal midline incisions parallel to and on either side of the spine[18]. The ovaries were located and dissected free from attachments[18]. All incisions were sutured, wound clipped and sterile glue applied (vetbond).

### 17-beta estradiol silastic capsule preparation and administration

KC mice were castrated or ovariectomized at age 6-7 weeks. The first 17-beta estradiol silastic capsule (or sham implant) was implanted 14 days later, and replaced every 4 weeks up to age 9 months. Silastic tubing (Silastic Laboratory Tubing, 1.58 mm inside diameter × 3.18 mm outside diameter, catalog no. 508-008, Dow Corning) was cut to 4.8mm. The tubing was sealed at one end with medical grade adhesive. The 4.8mm of the capsules was filled with 17-beta estradiol (E2) (17-beta estradiol, ≥99% pure, catalog no. 50-28-2). Silastic capsules were sealed with silastic medical adhesive, type A (product no. A-100, Dow Corning, purchased from Factor II). 17-beta Estradiol-filled silastic capsules have been shown to effectively increase estrogen levels in C57BL/6J mice when implanted as previously described[19,20]. Before implantation the capsules were soaked in sterile saline overnight at 37 °C. Mice were anesthetized with isoflurane for silastic capsule implantation and given slow-release buprenorphine (0.5 mg/kg sc). The back was shaved using clippers and sterilized with iodine scrub. An incision was made on the caudal aspect of the back just to the right of midline. Capsules were inserted parallel to the spine, and the incision was closed with wound clips.

### Statistical analysis

Comparisons of tumor development between groups was accomplished using the fisher’s exact test. The qPCR data was analyzed using an unpaired, two-tailed t-test with Welch’s correction to evaluate possible expression differences of *Pdx1* and *Cre* in the sample groups. Data was considered significant with a *p*-value <0.05.

## Results

### Development of anal tumors in KC mice

Of the 30 KC mice (16 KC males and 14 KC females) initially evaluated, 16.7% (5/30) developed pancreatic ductal adenocarcinoma (PDAC), which is consistent with previously published incidence in KC mice (S1A Fig) [8]. There were no statistically significant differences between development of PDAC precursor lesions (PanIN-1, PanIN-2, or PanIN-3) or PDAC in male and female KC mice (S1B Fig). Furthermore, 66.7% (20/30) of the KC mice possessed at least one external tumor on the body surface. Concordant with previous publications, roughly 36% of KC mice developed a facial lesion identified as facial papilloma [8,21] Notably, anal tumors were also observed and identified as invasive anal squamous cell carcinoma (anal SCC) on histopathologic analysis (Fig 1). The tumors became macroscopically visible after 5 months, were clearly evident by 6 months, and of significant size by 9 months (Fig 2). Mice with anal tumors displayed no increase in lethality, with normal mobility and weight gain up until the time of euthanasia (age 9 months). All anal SCC tumors were located at or just distal to the anorectal junction. All the tumors were characterized as grade 1. The neoplastic cells were well differentiated and easily recognized as squamous epithelium, infiltrating within a desmoplastic stroma with focal keratinization. Anal SCCs were localized to the anus, with no evidence of metastasis to distant organs. The pancreas, stomach, small intestine, colon, spleen, thymus, lungs and liver underwent gross analysis, but no evidence of metastasis from anal SCC tumors were found. Additionally, the stomach, spleen, pancreas, small intestine, and liver underwent histopathologic analysis with no evidence of metastatic spread. Only KC mice developed tumors (i.e. only mice possessing activated *Kras*-mutation), while age-matched male and female control mice did not develop external tumors (Table 1).

**Table 1.**
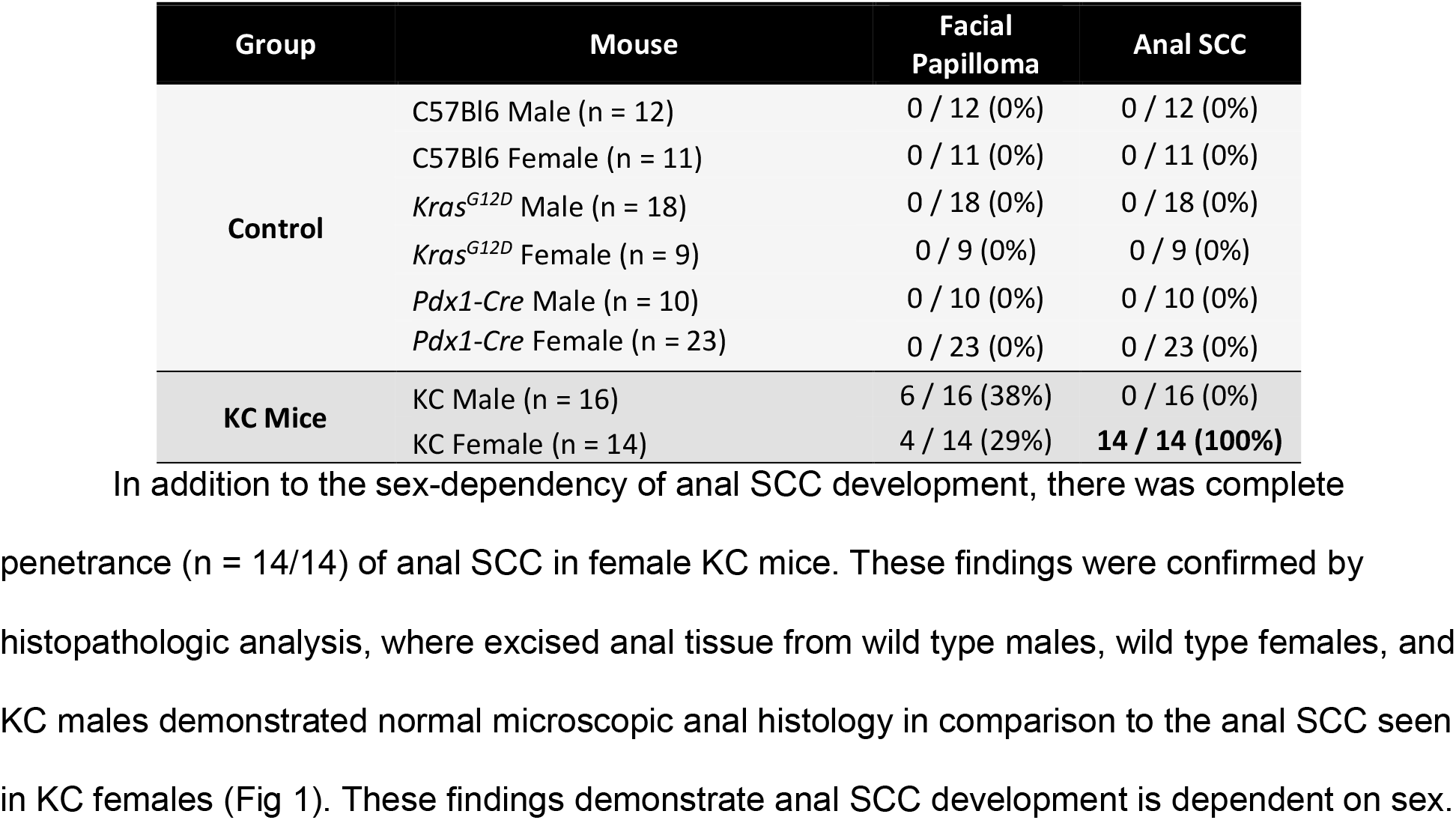
Sex-Dependent Incidence of Skin and Anal Lesions.

**Fig 1.**
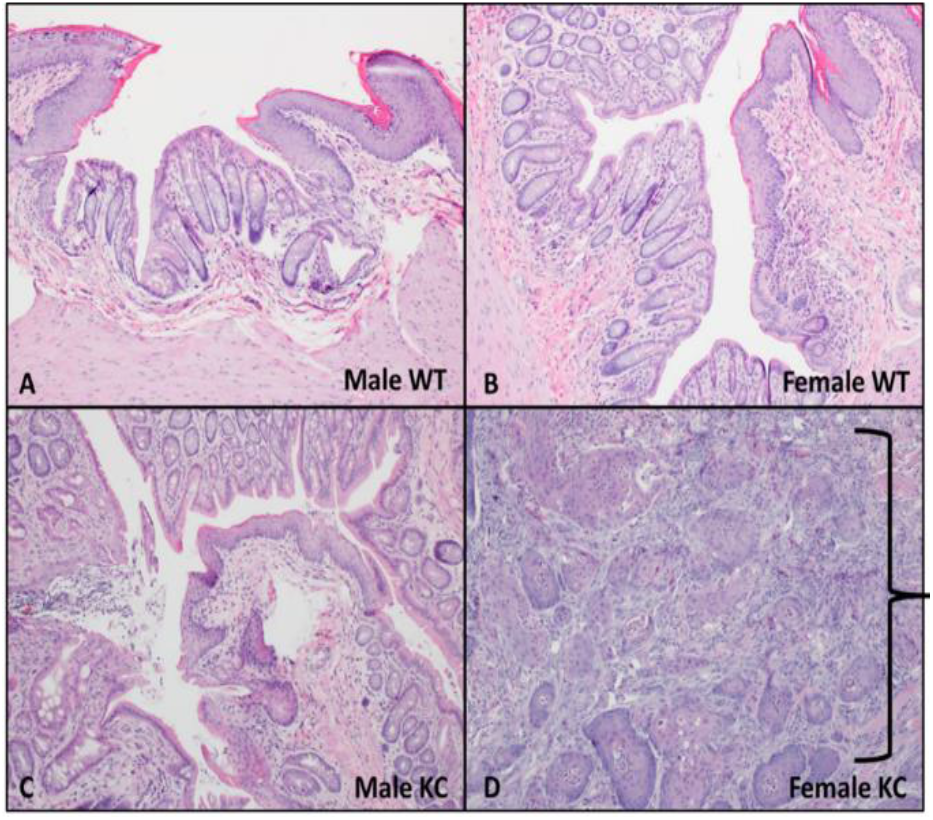
Anal tumor development in KC mice. At age 9 months, anorectal tissue was excised for histological analysis. In male (A) and female (B) C57Bl6 wild type mice, normal anorectal histology was present. Additionally, 9-month-old KC male mice (C) demonstrated normal anorectal histology. In contrast, large perianal tumors were grossly evident in 9-month-old female KC mice with invasive anal SCC present on histologic examination. This is indicated by the bracket (D).

**Fig 2.**
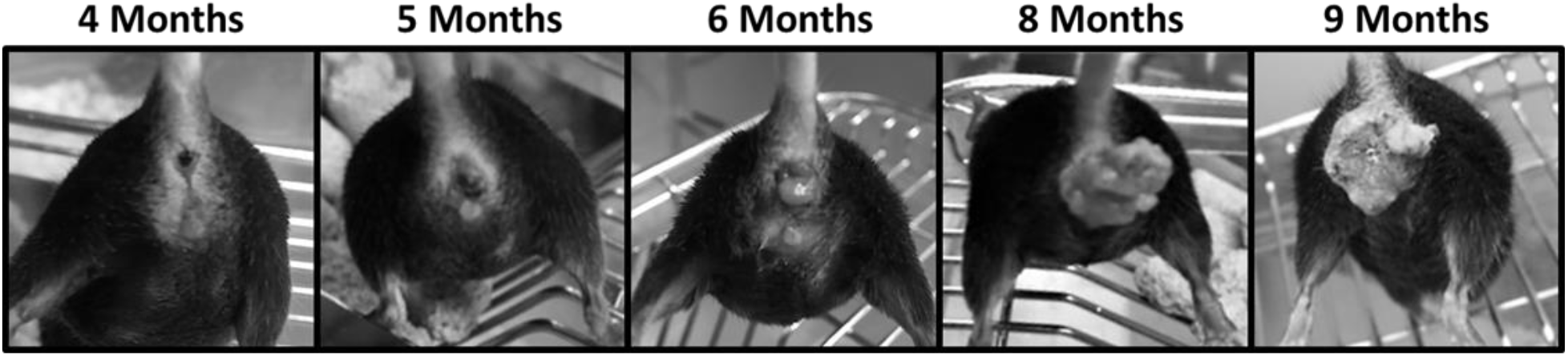
Anal tumor progression over time. In female KC mice, anal tumors are visible as an area of congestion at 4 months of age, with mild erythema around the anal region. By age 5 months, an anal tumor is generally evident. By 9 months of age, the anal tumors are significant in size. Despite the large size, the mice are able to maintain weight, consume food, and defecate normally. No mice experienced obstructive symptoms.

### Sex significantly influences anal SCC tumor development in KC mice

A significantly higher rate of external tumors was observed in female KC mice compared to male KC mice (100% vs 38%, p = 0.005). While there was no difference in the incidence of facial papilloma between the female and male KC mice (29% vs 38%, p = 0.65), there was a stark difference in anal SCC incidence, which occurred exclusively in female KC mice (100% vs 0%, p = 0.00001) (Table 1).

### Anal carcinogenesis in KC mice was not due to papillomavirus infection

Mouse papillomavirus (MmuPV1) has been associated with the development of anal disease and cancer in mice [14,22]. The animal facility where the mice are housed is routinely screened for MmuPV1 and it has not been detected in our colony, and immunocompetent C57Bl6 mice are known to rapidly clear MmuPV1 before tumor development occurs [22]. Furthermore, the anal tumors that developed in the KC mice were negative for characteristic features of papillomavirus-induced anal SCC during histopathologic evaluation. The overlying squamous mucosa did not exhibit koilocytosis, binucleation or raisanoid nuclei to suggest viral cytopathic effect from histopathology of KC mouse anus (Figure 3A) [23]. Representative anal SCC tumors in KC mice were evaluated for MmuPV1 viral transcripts using RNAScope and for MmuPV1 DNA using PCR.[14] No MmuPV1 signal was detected by RNAScope (Fig 3A), and no MmuPV1 DNA was detected within the anal tumors of the KC mice (Fig 3B). Together, these data show that the anal SCC in this study was not driven by papillomavirus infection.

**Fig 3.**
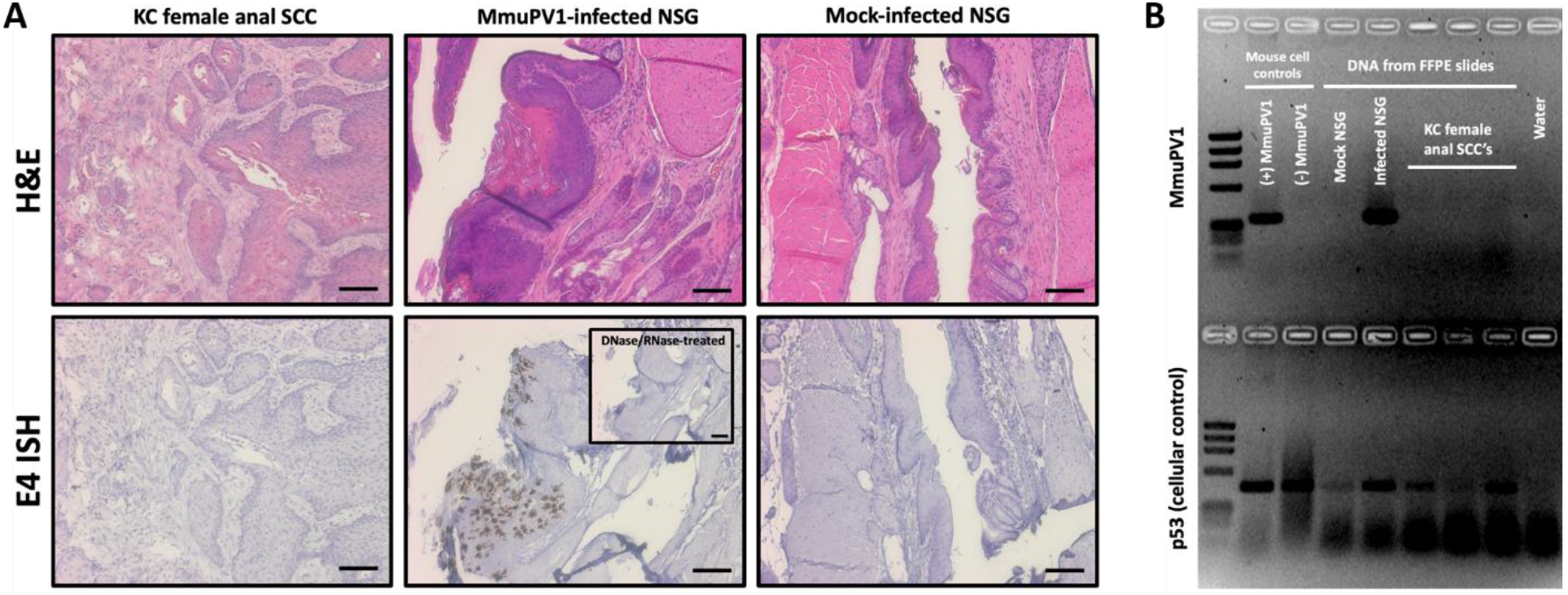
Anal squamous cell carcinomas in KC mice were negative for MmuPV1 infection. (A) No viral signatures were detected in representative anal tumors from KC mice stained via RNAScope ISH with a probe specific to the MmuPV1 E4 region of the genome. MmuPV1-infected and mock-infected Nod-*scid* IL2Rγ^null^ (NSG) mouse anal tissues were included as positive and negative controls, respectively. Scale bars equal 100 µm. (B) DNA was recovered from FFPE slides of representative KC female anal squamous cell carcinomas and MmuPV1-infected and mock-infected anal tissues, and PCR was performed using primers specific to the MmuPV1 genome. KC lesions were negative for MmuPV1 DNA.

### Kras^G12D^ mutation is present in anal tissue of KC mice

The absence of anal SCC in any control mice combined with the presence of anal tumors only in KC mice indicated *Kras*-mutation was likely responsible for the observed anal SCC. To confirm, we assessed anal tissue for expression of *Pdx1* mRNA, *Cre* mRNA, and activated *Kras*^*G12D*^ mutation (genomic DNA) in both male and female mice (Fig 4-6). In the KC model, the *Pdx1* promoter mediates the expression of *Cre* recombinase in both male and female anal tissues. We compared *Pdx1* mRNA levels in male and female WT and KC mouse anus at nine months of age (Fig 4A and 4B), and found similar levels of *Pdx1* expression amongst the WT male and female mice (Fig 4A). The presence of *Pdx1* in anal tissue has also been identified in prior investigations [24]. At nine months, expression of *Pdx1* in KC female anus / anal tumor was significantly higher than KC male anus (Fig 4B). We concordantly found *Cre*-recombinase expression was significantly higher in KC females compared to KC males (Fig 4C). WT mice did not express *Cre*-recombinase due to the absence of *Pdx1-Cre* transgene. The difference in *Pdx1* and *Cre* mRNA expression at nine months of age was likely related to evaluation of KC female anus / tumor (tumor tissue harboring more *Pdx1-Cre* expressing cells) versus non-tumor anal tissue of males. Thus, we assessed *Pdx1* and *Cre* expression at five months, prior to onset of macroscopic tumor and found no differences in levels of *Cre* and *Pdx1* expression between the groups of mice (Fig 4D-4F). To substantiate our expression data, we crossed the KC mice to Ai14 mice to generate the AiKC ‘marker’ mouse model (*Rosa26*^*LSL-tdTomato*^ ; *LSLKras*^*G12D*^ ; *Pdx1-Cre)*. AiKC mice harbor the LSL-tdTomato red fluorescent protein in the Rosa26 locus, and in the presence of Cre-recombinase, the stop sequence is excised allowing for expression of tdTomato protein, localizing Pdx1 and Cre expression and serving as a marker for expression of activated *Kras*^*G12D*^ gene. Both male and female AiKC mice (Fig 5) displayed tdTomato in the anal canal epithelium confirming Pdx1 and Cre expression and providing an expected localization for mutant *Kras*^*G12D*^ expression. Concordantly, isolation of genomic DNA from anal tissue of male and female KC mice demonstrated the activated *Kras*^*G12D*^ mutation (Fig 6 and S2 Fig), which was absent in WT mice. When excising the anu from female KC mice, the specimen was removed *en bloc* with the large anal tumor, and DNA isolation revealed clear presence of the activated *Kras*^*G12D*^ mutation (Fig 6, S2 Fig). The male KC anal tissue appeared grossly and histologically normal, yet genomic DNA isolated from whole anal tissue demonstrated the same activated *Kras*^*G12D*^ mutation (Fig 6, S2 Fig). As a follow-up, the anus from KC mice at the earliest timepoint allowable (age 6-8 weeks) was excised and genomic DNA isolated (S3 Fig). Activated *Kras*^*G12D*^-mutation was present in both sexes indicating early expression of the oncogene. This was concurrent with the clear evidence of tdTomato (Pdx1/Cre) expression at age 12 weeks in the male and female AiKC mice. Interestingly, we were unable to detect a mutant-*Kras* band (or only a faint band) in one male KC mice (Fig 6), which was likely due to detection error from the excised anal samples. Thus, to confirm, we used FFPE sections generated from the 9-month male KC mice (cohort used to evaluate for tumor formation) to specifically evaluate genomic DNA from the anal canal epithelium. These sections should harbor the activated *Kras*^*G12D*^ mutation based on results of tdTomato staining (AiKC mice), and notably, we found that all male mice express mutant *Kras*^*G12D*^ (S4 Fig). Together, these data demonstrate the presence of activated *Kras*^*G12D*^ mutation in both male and female KC anus, yet an absence of tumor formation in male KC mice.

**Fig 4.**
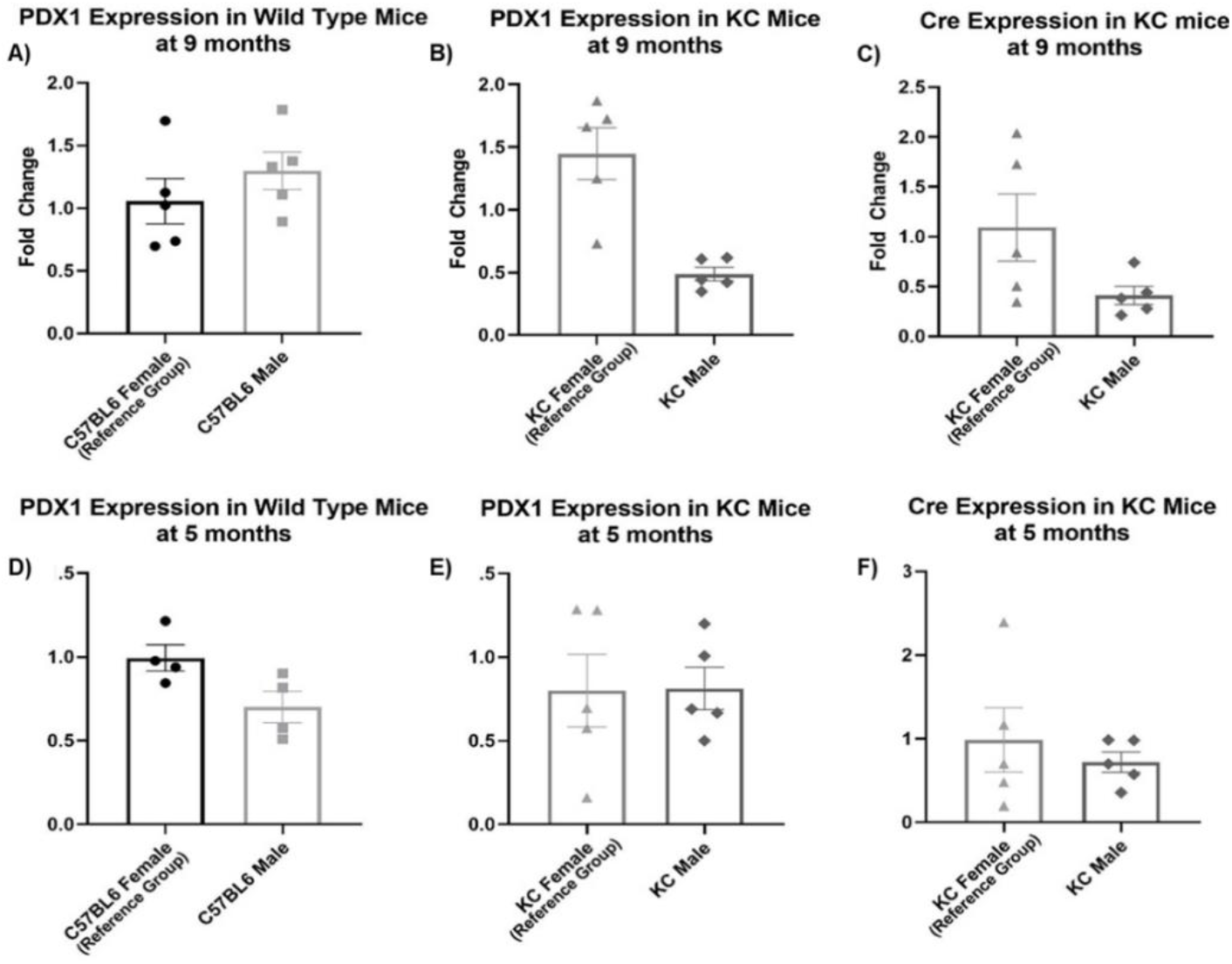
Pdx1 expression and Cre-recombinase expression in mouse anus. Anal tissue was excised from male and female wild type mice and male and female KC mice at 9 and 5 months of age. mRNA was isolated and RT-PCR performed to evaluate for *Pdx1* and *Cre*-recombinase expression. Male and female C57BL/6J WT mice demonstrate *Pdx1* expression in anal tissue at both 5 and 9 months, as did male and female KC mice (A,B,D,E). At 9 months age, KC females express significantly higher amounts of *Cre* mRNA (C) due to the presence of tumor tissue. At 5 months, male and female KC mice express similar amounts of *Pdx1* and *Cre* mRNA (E,F).

**Figure 5:**
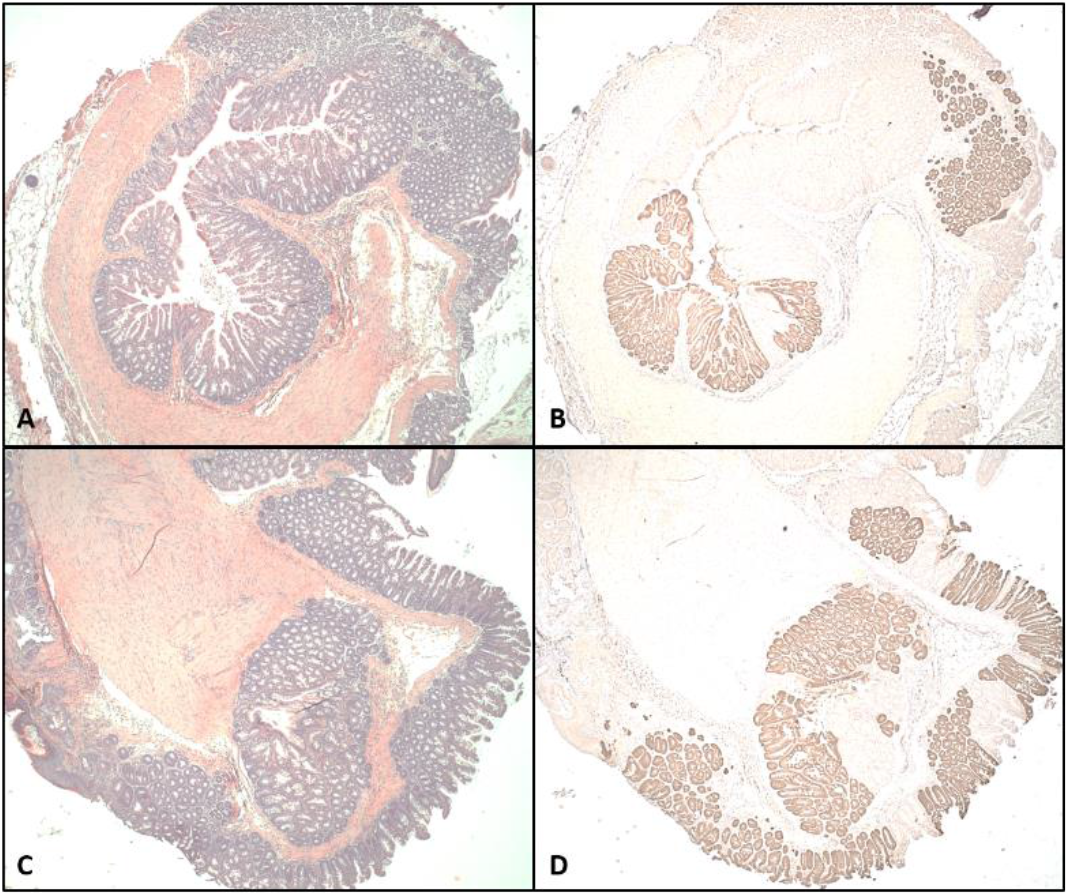
Tdtomato expression indicates Pdx1-Cre expression and mutant Kras expression in both male and female mice. Anal tissue was excised from 3 month old *Ai14* ; *LSL-Kras*^*G12D*^ ; *Pdx1-Cre (AiKC)* female (A, B) and male (C, D) mice, fixed and frozen embedded and sectioned for H&E analysis (A and C). Additional adjacent sections were prepared for immunohistochemistry (IHC) to identify tdTomato protein (B and D). Positive IHC signal for tdTomato reveals the location of Cre expression (*Pdx1-Cre*), which serves as a marker for the location of mutant *Kras*^*G12D*^ expression.

**Fig 6.**
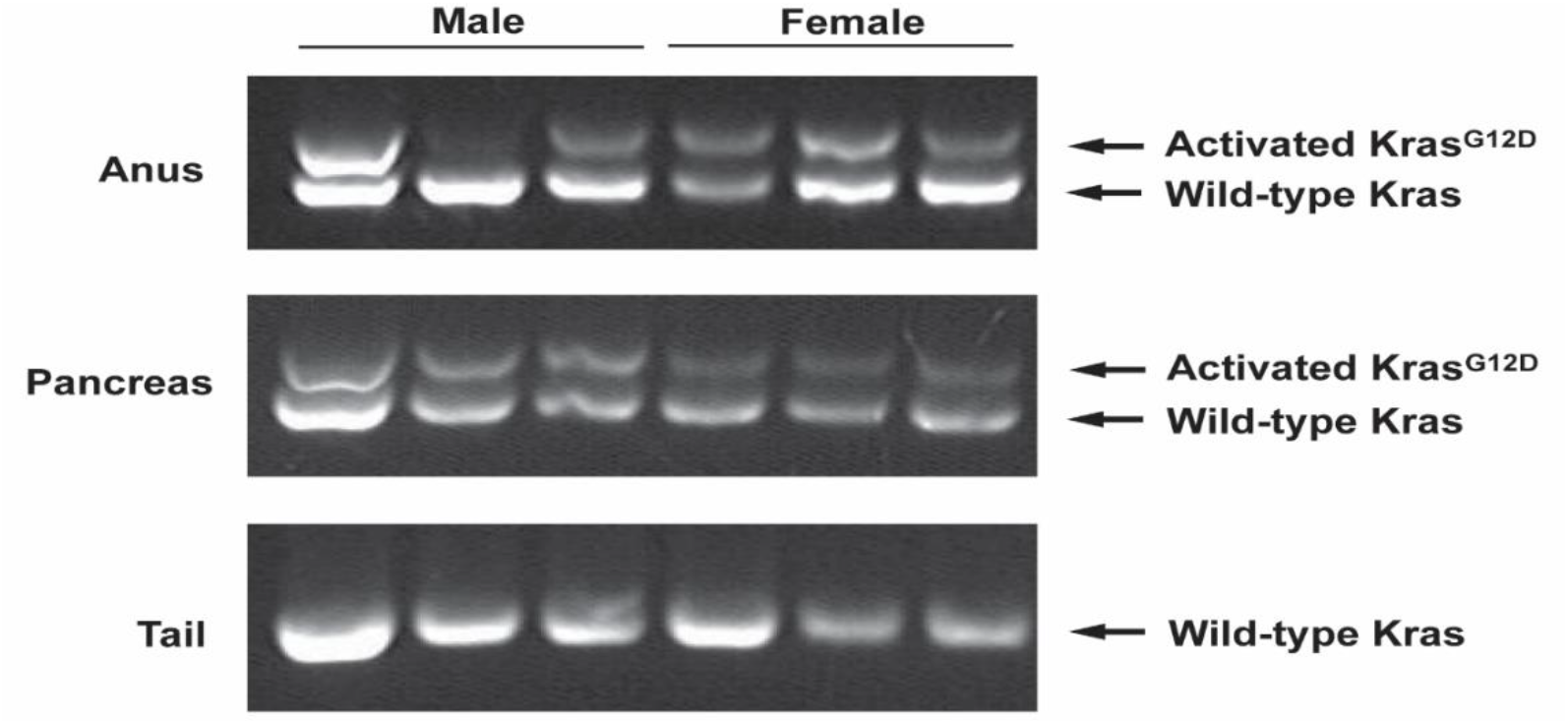
Activated *Kras*^*G12D*^ is present in the anal SCC anal tumor tissue. The activated *Kras*^*G12D*^ mutation is detected in the female anal tumor tissue, and the male anal tissues. Activated *Kras* refers to the successful removal of the *Lox-stop-Lox* codon preceding the *Kras*^*G12D*^. The activated mutation is present in the pancreas due to extensive *Pdx1* expression, and absent in tail samples, which lack *Pdx1* expression. Positive and negative controls are shown in supplemental figure 2.

### Sex hormone dependence of anal SCC formation

To discern why only female KC mice develop anal tumors despite the presence of activated *Kras*^*G12D*^ mutation in both male and female KC mice, we assessed sex hormone dependence. The roles of male and female sex hormones in the development of these tumors were judged by castration of male mice (n = 11) and ovariectomy of female mice (n = 13) at 6-7 weeks of age, according to standardized protocol.[17,18] Castration dramatically lowers the amount of testosterone that is produced in male mice[17] and, similarly, ovariectomy significantly lowers the amount of estrogen/progesterone produced in female mice.[18,25] Castrated KC males displayed an unchanged phenotype compared to intact KC males, with none of the mice developing anal SCC (0/11 KC castrated males vs 0/16 KC intact males, p = 1) (Table 2). In contrast, ovariectomized KC female mice exhibited a striking change compared to intact KC female mice, with only 15% of the ovariectomized cohort developing macroscopic anal lesions (2/13 ovariectomized KC females vs 14/14 intact KC females (P<0.0001)) (Table 2). This was confirmed on microscopic analysis, in which the eleven ovariectomized KC female mice without macroscopic tumors demonstrated normal anal histology (i.e. no microscopic tumors or dysplasia). (Figure 7). This remarkable finding indicates that female sex hormones are crucial for *Kras*^*G12D*^-driven anal SCC development in KC mice.

**Table 2:** Sex-Dependent Incidence of Skin and Anal Lesions after Castration or Ovariectomy

| Group | Mouse | Facial Papilloma | Anal SCC |
| --- | --- | --- | --- |
| **Castrated/Ovariectomized Controls** | C57Bl6 Male (n = 2) | 0 / 2 (0%) | 0 / 2 (0%) |
|  | C57Bl6 Female (n = 2) | 0 / 2 (0%) | 0 / 2 (0%) |
|  | *Kras*<sup>G12D</sup> Male (n = 2) | 0 / 2 (0%) | 0 / 2 (0%) |
|  | *Kras*<sup>G12D</sup> Female (n = 2) | 0 / 2 (0%) | 0 / 2 (0%) |
|  | Pdx1-Cre Male (n = 2) | 0 / 2 (0%) | 0 / 2 (0%) |
|  | Pdx1-Cre Female (n = 2) | 0 / 2 (0%) | 0 / 2 (0%) |
| **Castrated/Ovariectomized KC Mice** | Castrated KC Male (n = 11) | 4 / 11 (33%) | 0 / 11 (0%) |
|  | Ovariectomized KC Female (n = 13) | 4 / 13 (30%) | 2 / 13 (15%) |

**Fig 7.**
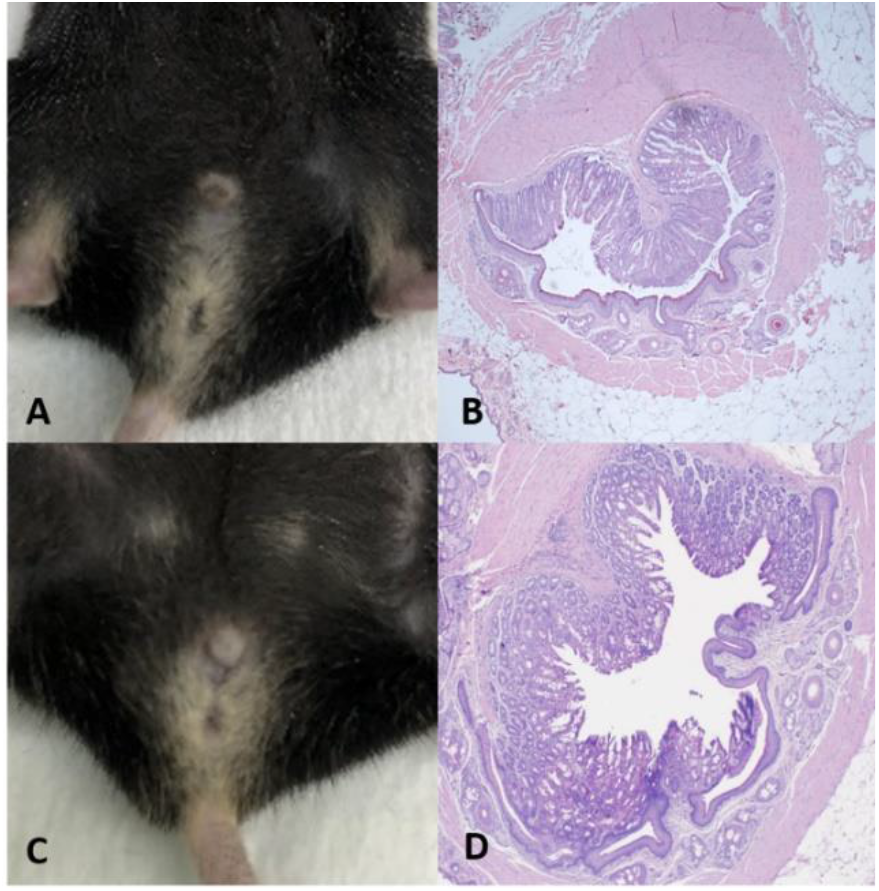
Castrated and ovariectomized KC mice and assessment for anal tumor. Male KC mice were castrated and female KC mice were ovariectomized at 6-7 weeks of age, and the anal tissue evaluated at age 9 months. Macroscopic examination revealed no abnormalities in castrated KC males and most ovariectomized KC females (representative images, A and C). Concordantly, histopathologic examination revealed no microscopic tumor formation in male KC mice or macroscopically normal ovariectomized KC female mice (representative images B and D, 40x) (D).

### Estrogen dependence of anal SCC formation

Estrogen has a known role in the development of several tumors [26–31] and a correlation with *Kras*-mutant cancer [28–31]. Thus, we tested whether the anal SCC tumor development in a papillomavirus-negative context was driven by estrogen. Ovariectomized and castrated KC mice were dosed with physiologic levels of 17-β-estradiol (N=5 for ovariectomized and N=4 for castrated) or sham dosed as a control (N=6 for ovariectomized and N=5 for castrated). To confirm successful E2 administration, uterine weights were assessed (S5 Fig).[20] E2 dosed females should remain in proestrus and thus have normal uterine weights while the sham dosed mice will have significantly lower uterine weights (S5A Fig) [20]. This standard approach enables accurate determination of estrogen reduction as opposed to a single timepoint (serum) which can vary substantially even in wild type (intact) mice.[20] Sham dosed mice demonstrated significant decrease in uterine weights confirming successful reduction in estrogen levels, while all E2 dosed female KC mice possessed normal uterine weights (i.e. intact) revealing appropriate and sufficient exogenous estrogen administration (S5A Fig). In the sham-dosed ovariectomized KC female group only 33% (2/6) developed a tumor (Table 3), concordant with results seen in the untreated ovariectomized mice (33% vs 15%, p-value =0.56) (Table 2). Meanwhile, in the beta-estradiol (E2) dosed ovariectomized KC female mice, 100% (5/5) developed macroscopically visible anal tumors by 4 months of age (Table 3), ‘rescuing’ the tumor phenotype and again demonstrating stark contrast to ovariectomized mice (100% vs 15%, p-value = 0.001). In the sham-dosed castrated KC male group, 0% (0/5) developed a tumor (Table 3), consistent with the results seen in the untreated castrated KC male mice (0/5 vs 0/11, p-value = 1). In contrast, 75% (3/4) of E2-dosed castrated KC male mice developed anal SCC that was macroscopically visible by 8 months of age and confirmed on histopathologic analysis (Fig 8). This remarkable and significant increase in tumor formation in E2-dosed KC males (75% vs 0%, p-value = 0.0088), when coalesced with the novel data from KC females, demonstrates an estrogen mediated sex-dependent development of *Kras*-mutant anal SCC.

**Table 3:** Incidence of Anal SCC in beta-estradiol dosed KC mice.

| Group | Mouse | Anal SCC |
| --- | --- | --- |
| Intact KC Mice | Intact KC Male (n = 16) | 0 / 16 (0%) |
|  | Intact KC Female (n = 14) | 14 / 14 (100%) |
| Castrated/Ovariectomized KC Mice | Castrated KC Male (n = 11) | 0 / 11 (0%) |
|  | Ovariectomized KC Female (n = 13) | 2 / 13 (15%) |
| Dosed and Castrated/Ovariectomized KC Mice | Sham Dosed Castrated KC Male (n = 5) | 0 / 5 (0%) |
|  | Beta-Estradiol Dosed Castrated KC Male (n = 4) | 3 / 4 (75%) |
|  | Sham Dosed Ovariectomized KC Female (n = 6) | 2 / 6 (33%) |
|  | Beta-Estradiol Dosed Ovariectomized KC Female (n = 6) | 5 / 5 (100%) |

**Fig 8.**
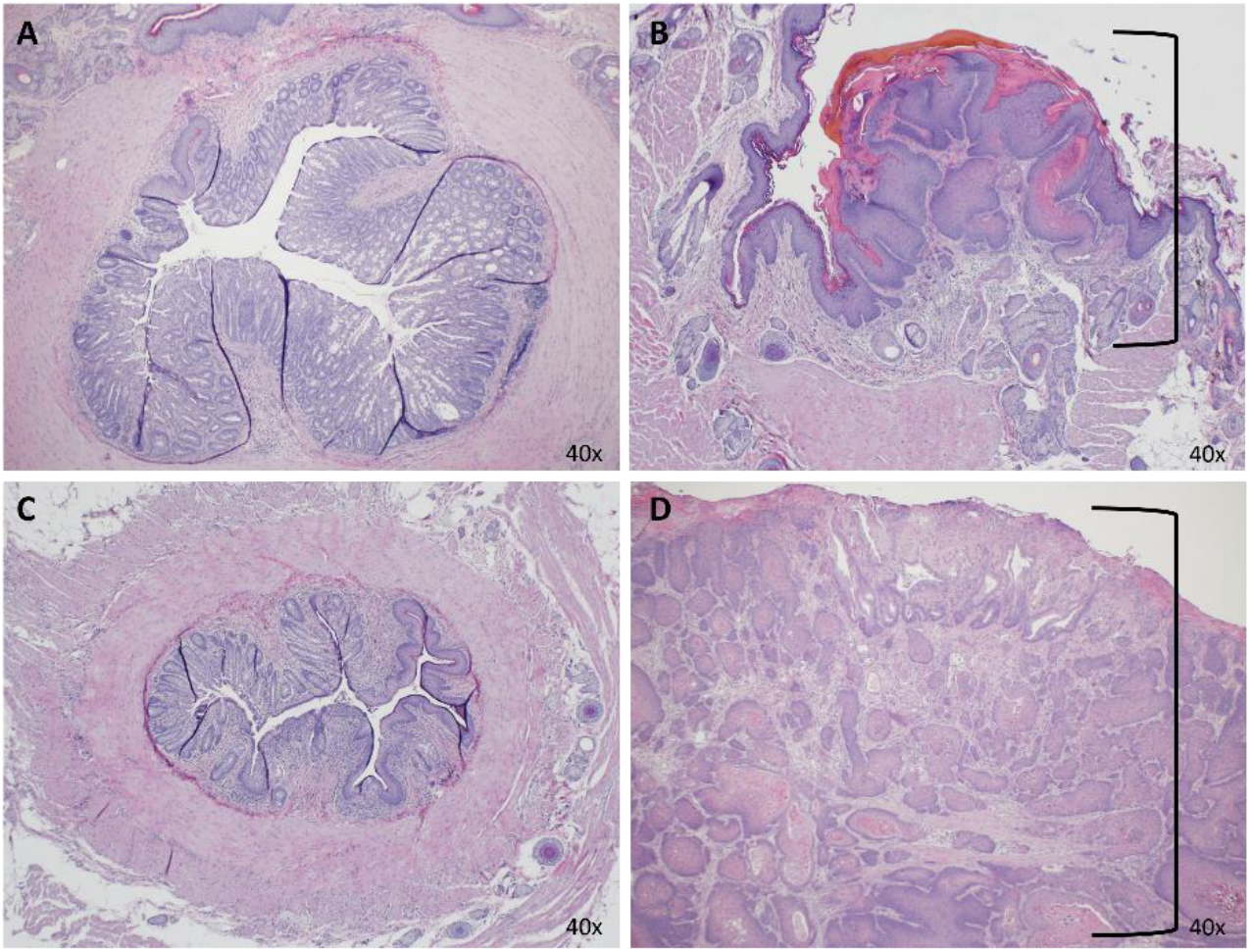
Estrogen dosed castrated KC male develop anal SCC. Castrated KC male mice and ovariectomized KC female mice were treated with 17-beta estradiol (E2) or sham (sesame oil). Sham dosed KC males did not develop anal SCC (A), while E2 dosing induced anal SCC in KC males (B, shown by bracket). Similarly, sham dosed KC females did not develop anal SCC (C) while E2 dosing rescued the anal SCC phenotype in KC females (D, shown by bracket).

### Estrogen Receptor present in male and female anus and female anal tumors

Estrogen signaling is mediated through two distinct receptors, ERα and ERβ.[32] Estrogen signaling through ERα has been shown to increase cellular proliferation, particularly within the mammary gland and uterus, while ERβ has been shown to counteract the proliferative effects of ERα.[32] Given the proliferation-inducing role of ERα, we expected that the anal epithelium and anal tumors would display ERα expression. To investigate, we performed fluorescent IHC using an antibody specific for ERα [15] in the male and female anus. In particular, we assessed ERα in intact KC females and found robust presence of ERα in the anal tumor (Fig 9). Moreover, we analyzed normal anus of ovariectomized KC females (no tumor) and KC males and again identified strong presence of ERα in the anal tissue, indicating exogenous estrogen in male (E2 dosed) and endogenous estrogen in female KC mice can bind receptor to induce tumor formation (i.e. receptor is not just expressed in tumor). Together, this substantive data suggests that estrogen binding to ERα potentiates mutant-*Kras*^*G12D*^ induced development of anal SCC in KC mice. (Fig 9).

**Fig 9.**
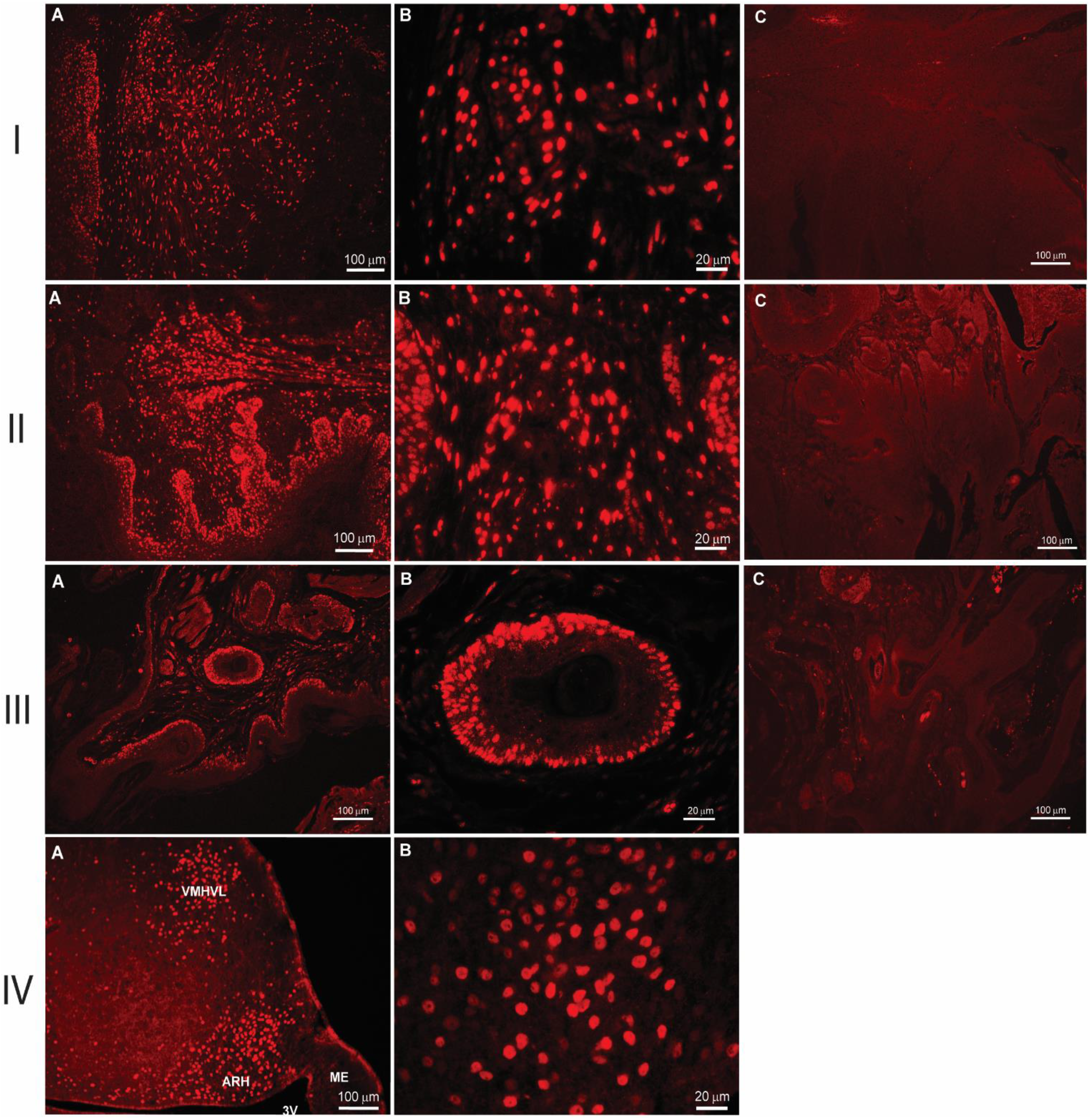
Estrogen Receptor alpha (ERα) is present in the KC anal tissue. The presence of ERα was detected using immunofluorescent IHC. ERα was found to be present in the anal SCC of intact KC female mice (I), in the anal epithelium of ovariectomized KC female anus lacking tumor formation (II) and in the KC male anus (III). Panels A and B show the fluorescent staining of the receptor at 10x and 40x respectively. Panel C in each group shows the tissue with only the use of the secondary antibody confirming no off target staining. Group IV shows ERα staining of mouse arcuate nucleus as the positive control.[15]

## Discussion

In this study, we identified that *LSL-Kras*^*G12D*^; *Pdx1-Cre* (KC) mice showed female-specific anal SCC development. Although this is a highly utilized genetically engineered mouse model in the study of PDAC, this novel phenotype has likely been overlooked for several reasons. While earlier studies of KC mice included only males, more recent investigations have included both male and female *Kras*-mutant mice to assess the development of PDAC. However, these studies focused on concomitant genetic mutations (e.g., *Trp53, Ink4a/Arf*)[9,33] or the influence of environmental changes (e.g., high-fat diet)[34] in addition to the *Pdx1*-*Cre* driven *Kras*^*G12D*^-mutation, which facilitate the onset of PDAC and consequent death at an early age (roughly four months for *LSL-Kras*^*G12D/+*^ ; *Trp53*^*-/-*^ ; *Pdx1-Cre* mice in our laboratory). Thus, because these tumors were only identifiable starting at 5-6 months age, studies with combination genetic mutations or environmental changes that caused earlier evaluation / demise in male and female KC mice may have conceivably missed the onset of anal SCC growth in female mice.

Approximately 85-95% of anal SCC development in humans is due to HPV infection, while the remaining 5-15% of cases occur from an unknown etiology.[4] Although mice can be infected in the anal tract with the mouse papillomavirus MmuPV1 [14,22], the KC mice in this study did not develop anal SCC as a result of MmuPV1 or HPV infection. Immunocompetent mice, such as the KC mice, have been shown to rapidly clear MmuPV1 in the anal tract.[22] Furthermore, there was no evidence of papillomavirus in the colony of this study, on histopathologic analysis, using RNA Scope or via PCR analysis of the anal SCC that developed within the KC mice. This is an important finding when considering that roughly 5-15% of anal SCC patients do not harbor papillomavirus as the underlying etiology and suggests this is a potential model of interest in studying the etiology of non-papillomavirus induced anal SCC.

In this study, the development of anal SCC in the KC mice is due to the activation of the *Kras*^*G12D*^ mutation in the anal tissue, which subsequently drives tumor formation. In the KC mouse model, the activation of the *Kras*^*G12D*^ mutation is controlled by cells expressing Cre-recombinase from the *Pdx-1* promoter.[8] Although only female KC mice develop anal SCC, both sexes of KC mice were shown to have equivalent expression of *Pdx1* and *Cre* in the anal epithelium at age 5 months. Furthermore, the expression tdTomato was clearly evident in both male and female AiKC mouse anus, which reveals the location of Pdx1-mediated excision of lox-stop and consequent tdTomato expression (surrogate for location of *Kras*-mutation). Along with the observation that only KC mice formed anal SCC, this data shows that the development of anal SCC is driven by the expression of Pdx-1 and Cre in the anal tissue resulting in activation of the mutated *Kras*^*G12D*^ gene. While we are not the first group to describe *Pdx1* expression outside the pancreas leading to the development of SCC tumors [24], to our knowledge, we are the first group to identify both *Pdx1* and activated *Kras*-mutation in anal tissue of KC mice and resultant anal SCC formation. Furthermore, the evidence presented disputes the possibility that sex-bias differences in anal tumor formation was simply due to the absence of *Pdx1*, Cre or activated *Kras*^*G12D*^ expression in male anal tissue.

Notably, the activated *Kras*-mutation did not appear to be present (or very faintly present) in all male anal samples retrieved by whole-excision and analyzed by PCR (one sample of the 9 month cohort). This was likely due to detection error when isolating DNA from whole male anus (i.e. hair follicles, glands, skin, colon), which prompted PCR analysis of genomic DNA from FFPE anus samples (containing anal epithelium) to confirm the presence of activated *Kras*-mutation in KC male anus.

To understand the female sex predilection for development of anal SCC, we evaluated the roles of sex hormones, as these are likely candidates contributing towards the observed phenotype. We castrated male mice to achieve significant reduction in circulating testosterone and ovariectomized female mice to reduce systemic production of female sex hormones. This standard approach is the most accurate method to determine sex-hormone dependence.[17–20] Castration did not alter the anal phenotype of male KC mice suggesting that the lack of testosterone does not modify the development of anal SCC. In contrast, only two out of thirteen ovariectomized mice developed anal SCC suggesting the tumor development was almost entirely dependent on female physiologic levels of estrogen/progesterone.

To further evaluate the involvement of the female sex hormones, we dosed ovariectomized KC female mice and castrated KC male mice with estrogen to see if this would result in the tumor phenotype. We chose to evaluate estrogen because of the strong correlation with other *Kras*-driven cancers. For example, a study done by Hammond et al. [28] used *LSL-Kras*^*G12D*^ mice (K mice in that study) to investigate the sex-differences seen in the development of lung adenocarcinomas. The methodology in this study included ovariectomy in female K mice followed by activation of the *Kras*^*G12D*^ -mutation through intra-nasal exposure of an adenoviral vector expressing Cre recombinase (AdeCre). The authors found a significant reduction in lung tumor burden (quantity and size) compared to intact females. Concordant with the current study, they successfully rescued the phenotype through estrogen administration using silastic capsules.[28] Furthermore, studies have shown estrogen mediates the development of mutant-*Kras*-driven endometrial cancer, ovarian cancer and vaginal SCC.[29–31] It has been shown that ERα is present in 4% of human anal SCC samples[26] and that estrogen is essential for activating cell proliferation of human epithelial SCC cell lines.[35] Thus, from these data, it is feasible that estrogens can ultimately influence *Kras*-induced non-papillomavirus anal SCC development, resulting in the sex-dependent development of anal SCC phenotype in female KC mice. Notably, in our study, ovariectomized KC mice that were dosed with physiologic levels of beta-estradiol (E2) developed anal SCC at 4 months of age, ‘rescuing’ the tumor phenotype. Furthermore, E2 dosed castrated KC male mice (equivalent dose as females) also developed anal SCC, albeit with a relatively delayed macroscopic onset (8 months) compared to E2 dosed females. This delayed onset despite equivalent E2 dosing may be due to differences in the number of ERα expressing cells or amount of ERα present in anal tissue, and will be evaluated in future studies. Regardless, the data presented strongly suggests that the development of anal SCC in KC mice is *Kras*-driven and estrogen mediated.

It is important to note that following reduction of estrogen (ovariectomy), 15% (2/13) of female KC mice and 2/6 (33%) sham dosed female KC mice still developed tumors. To confirm that our ovariectomized KC female mice did experienced a significant reduction in circulating estrogen, and that the E2 dosed mice possessed sufficient levels of circulating estrogen, we used a standardized approach of uterine weights. This methodology is more accurate than ‘single time point’ levels of estrogen in circulating blood, due to the substantial variation of circulating estrogen in normal females.[20] In contrast, uterine weights reflect the steady levels of estrogen stimulation over an extended period. These techniques helped to confirm successful reduction of (ovariectomy) and rescue of (E2 dosed) circulating estrogen. Although our study revealed a clear correlation between estrogen and tumor formation, and a dramatic change in tumor phenotype with reduction of estrogen, it remains unclear why few ovariectomized KC mice still developed tumors; this may have been related to tumor initiation prior to the onset of ovariectomy. In follow up studies, we will use ERα knock out mice (B6.129P2-*Esr1*^*tm1Ksk*^/J) crossed to KC mice to investigate whether innate absence of estrogen to bind ERα prevents tumor formation. It is also currently unknown how estrogen and mutant-*Kras* synergistically drive anal tumor formation. Interestingly, one group has developed a non-HPV model of anal carcinogenesis, using tamoxifen-inducible deletions of *Pten* and *Tgfbr1*.[36] The authors found that anal SCC development was contingent upon STAT3 activation.[36] Cancers in other organ systems (e.g. pancreas cancer, lung cancer) have shown an interdependence between *Kras*^*G12D*^ mutation and heightened STAT3 activity[37,38], and it has also been shown that estradiol functions to increase STAT3 activation.[39] For example, estradiol was shown to increase STAT3 activation in female rat brain which results in neuroprotection against ischemic brain injury.[39] The association between estrogen and STAT3 activation along with the association between STAT3 activity and mutant *Kras*^*G12D*^-induced cancer formation suggests a possible mechanism behind the phenotype of sex-dependent anal SCC development in KC mice. Subsequent analyses will aim to clarify these questions and study limitations, as well as focus on elucidating the specific underlying mechanism by which estrogen enhances *Kras*-mutant anal SCC development.

Our study clearly shows the sex-dependent development of anal SCC is tied to presence of physiologic levels of estrogen in female mice and characterizes a clinically relevant subtype of anal SCC. The finding that the *Kras*-mutation is largely dependent upon estrogen to induce tumor development is an exciting result that may have direct clinical applicability for patients with non-HPV anal SCC that have poorly understood pathogenesis and are known to exhibit resistance to standard of care therapy.[5] Additionally, with the previously unidentified observation of *Pdx1*-driven *Kras*-mutation present in anal tissue of KC mice, the novel phenotype described in this study may also provide a new mouse model for evaluation of the non-papillomavirus subtype of anal SCC.

## Acknowledgments

This work was supported by the American Cancer Society (grant number IRG-15-213-51). The authors would like to thank the Experimental Animal Pathology Lab (EAPL) supported by the UWCCC (P30 CA014520) for use of its facilities and services. The authors would also like to thank Dr. Jing Zhang for providing positive control DNA for activated *Kras*-mutation from the bone marrow of *LSL Kras/+; Mx-1 cre/+* mice, Dr. Paul Lambert for input regarding MmuPV1-mediated anal tumor formation (funding CA210807), and Dr. Chad Vezina for insight into investigating sex-hormone driven cancers. We also thank Ms. Martha A. Bosch for technical assistance.

## Supporting Information

**S1 Fig.**
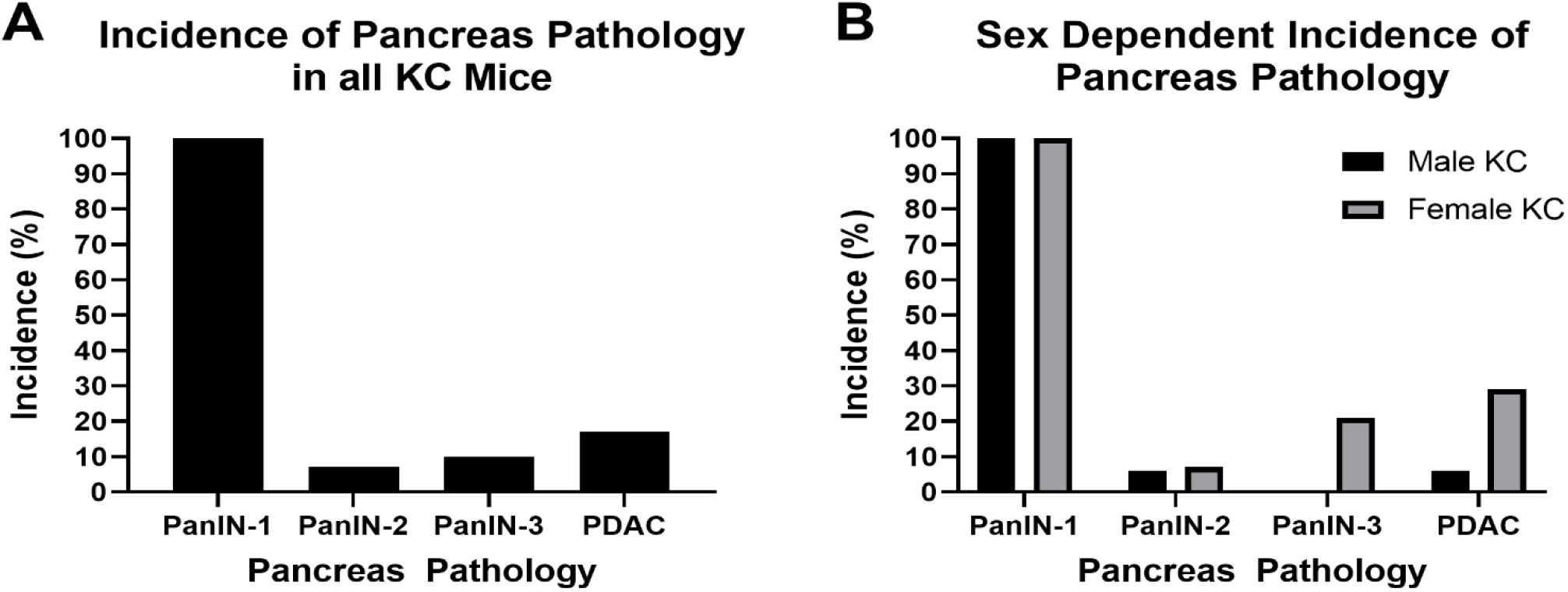
The incidence of pancreas pathology of male and female mice. A) The incidence of pancreatic precursor lesions (PanIN-1, PanIN-2, PanIN-3) and PDAC in all the KC mice match what was previously reported in this model. B) There is no statistically significant differences in development of pancreatic pathology between male(n= 16) and female KC (n – 14) mice. Both male and female show a 100% incidence of PanIN 1(p-value = 1). For PanIN1, PanIN 2, PanIN 3 and PDAC male vs female incidence with their p-vales are as follows: 100% vs 100% (p = 1), 6.25% vs 7.15% (P > 0.99), 0% vs 21.43% (P = 0.09) and 6.25% vs 28.57% (P = 0.16)

**S2 Fig.**
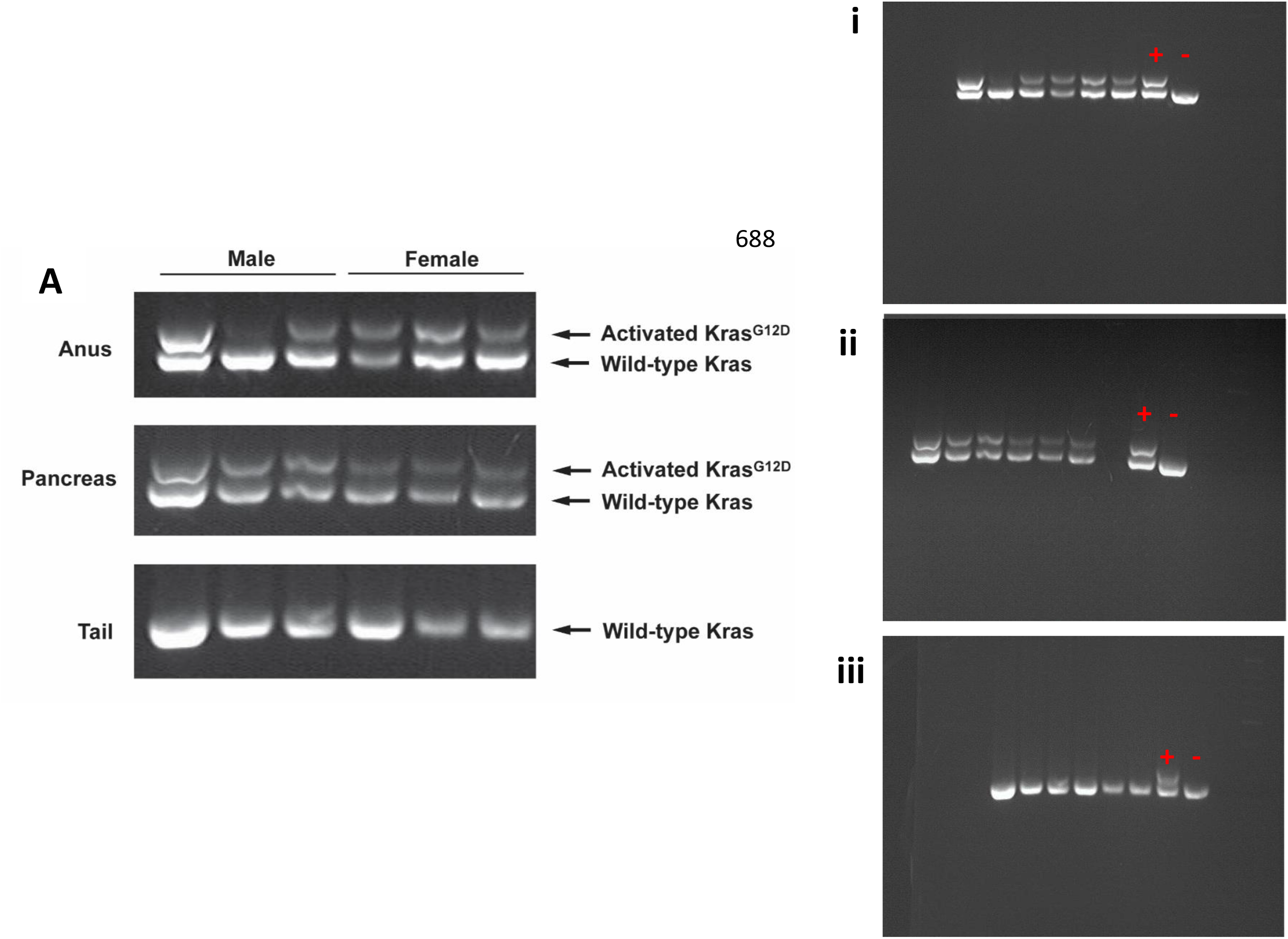
Genotyping for mutated Kras gene in 9 month old male and female KC mice. Full gel images of mutated Kras Genotyping in 9 month old male and female anus (i), pancreas (ii) and tail (iii) next to the cropped image used in the manuscript text (A). The ‘+’ and ‘-’ indicate the positive and negative control bands on each gel. The positive control is DNA from *LSL Kras/+; Mx-1 cre/+* mouse bone marrow containing the activated *Kras* gene (Control (+)) and the negative control is DNA from the pancreas of a C57BL6/J mice (Control (-)).

**S3 Fig.**
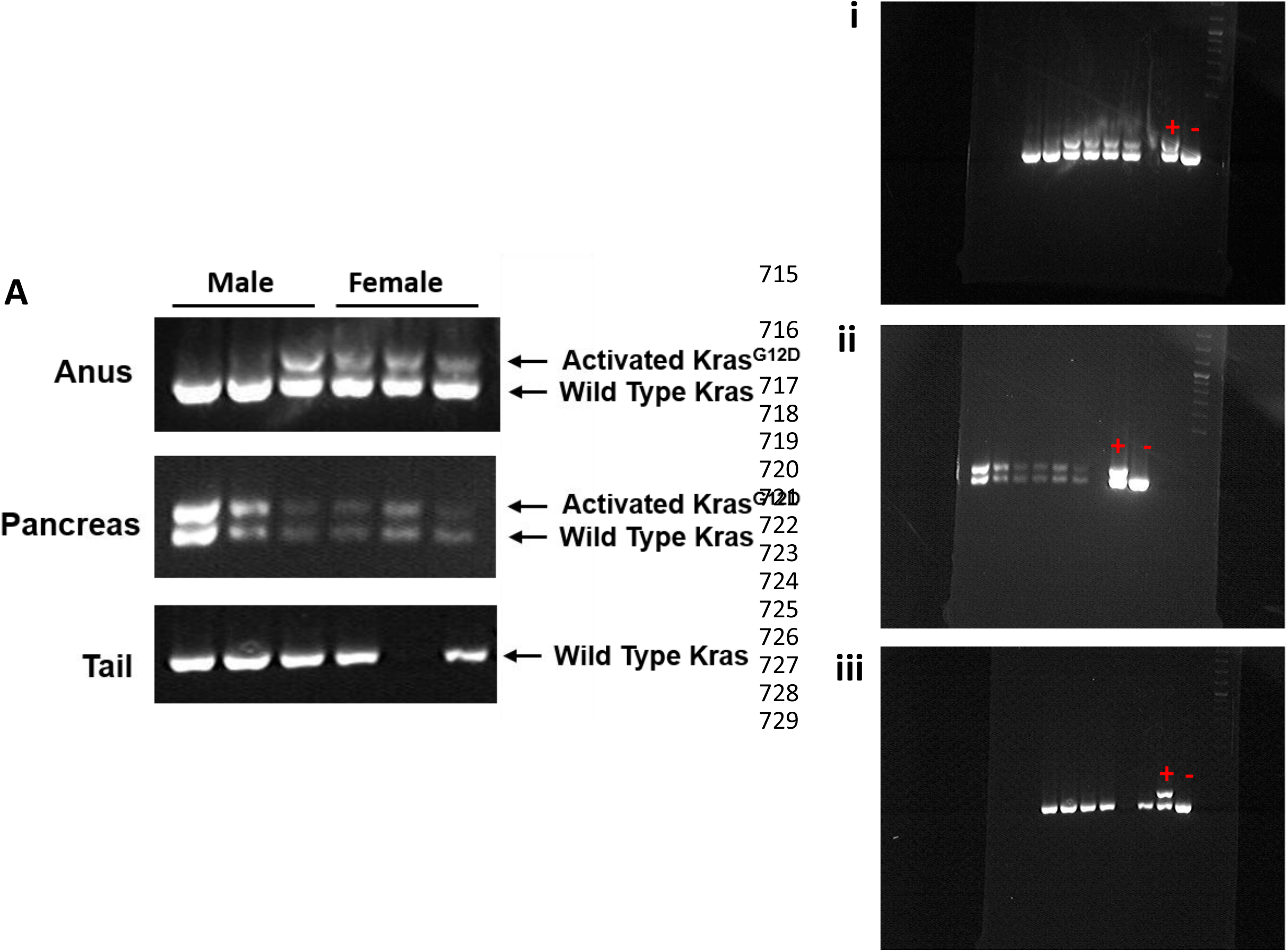
Genotyping for mutated Kras gene in 8 week old male and female KC mice. **A)** Full gel images taken of the PCR product. This shows that there are male KC anus’s with the mutated kras gene activated at an early age (8 weeks old). Despite the active mutated kras gene being present, no males develop anal SCC. The full gel images for the anus (i), pancreas (ii) and tail (iii) are to the right of the full image. The ‘+’ and ‘-’ indicate the positive and negative control bands on each gel. The positive control is DNA from *LSL Kras/+; Mx-1 cre/+* mouse bone marrow containing the activated *Kras* gene (Control (+)) and the negative control is DNA from the pancreas of a C57BL6/J mice (Control (-)).

**S4 Fig.**
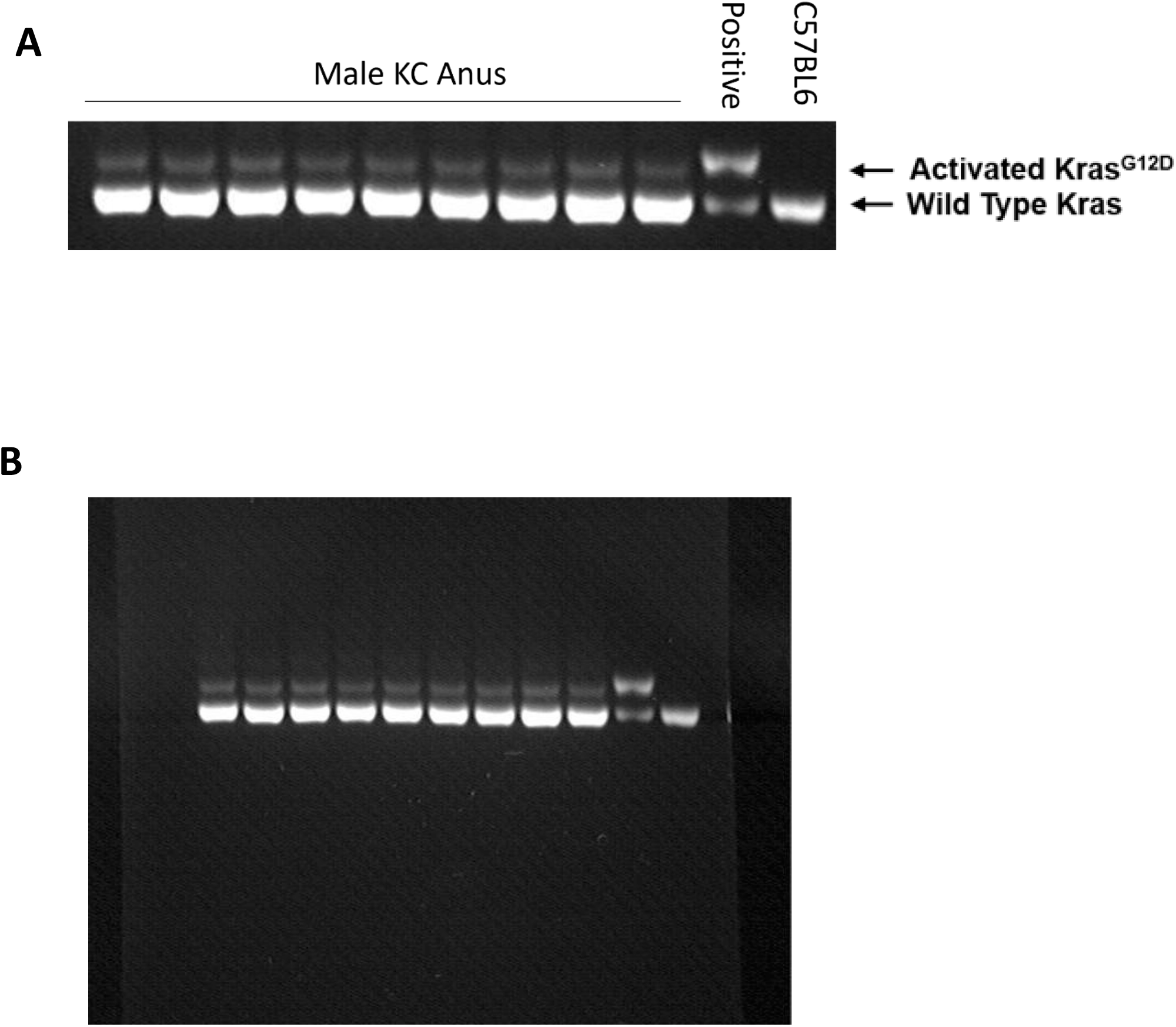
Anus from male KC mice display the activated *Kras* mutation. Genomic DNA was isolated from the available FFPE samples from age 9 month male KC mice and analyzed using PCR for the activated *Kras* mutation. All male anus showed the presence of the activated *Kras* mutation within the anal tissue (A). The full gel is shown in panel B.

**S5 Fig.**
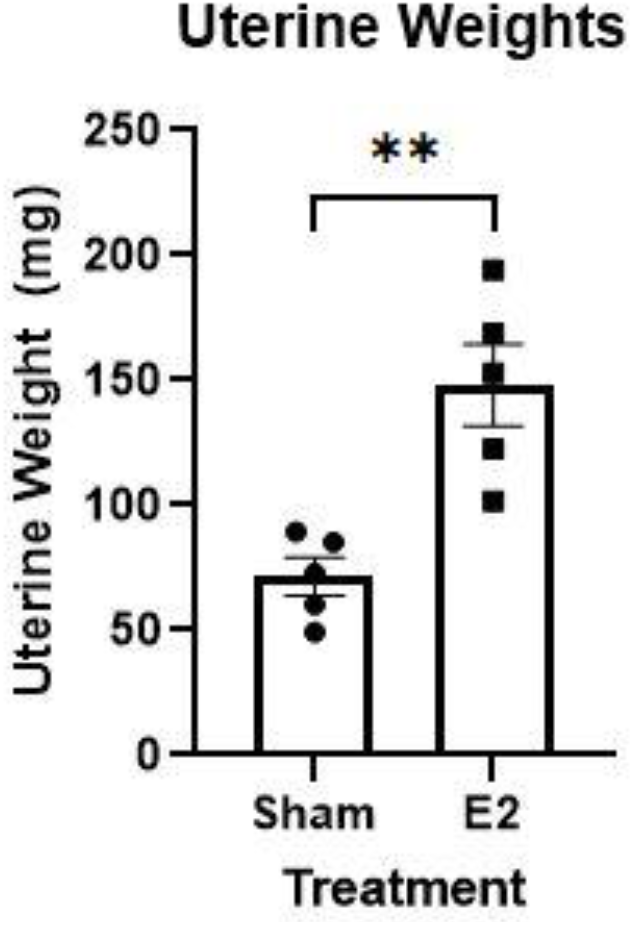
E2 dosed females show increased Uterine weights consistent with E2 dosing. The E2 dosed female mice have a significantly increased uterine weight compared to the sham dosed mice indicating successful E2 administration.

